# Characterizing the microbiome of patients with myeloproliferative neoplasms during a Mediterranean diet intervention

**DOI:** 10.1101/2023.01.25.525620

**Authors:** Julio Avelar-Barragan, Laura F. Mendez Luque, Jenny Nguyen, Hellen Nguyen, Andrew O. Odegaard, Angela G. Fleischman, Katrine L. Whiteson

## Abstract

Myeloproliferative neoplasms (MPN) are a class of hematological malignancies which result in the overproduction of myeloid lineage cells. These malignancies result in increased cytokine production and inflammation, which correlate with worsened symptom burden and prognosis. Other than bone marrow transplantation, there is no cure for myeloproliferative neoplasms. As such, treatments focus on reducing thrombotic risk, inflammation, and symptom burden. Because current pharmacological treatments carry significant side-effects, there is a need to explore low-risk therapies. One alternative is the Mediterranean diet, which is rich in anti-inflammatory foods, reduces inflammatory biomarkers, and beneficially alters the gut microbiome. Here, we performed a 15-week clinical trial of 28 individuals with MPN who were randomized to dietary counseling based on either a Mediterranean diet or standard U.S. Guidelines for Americans. Our primary objective was to determine if MPN patients were able to adopt a Mediterranean eating style when supported with dietician counseling. As exploratory endpoints, we investigated the impact of diet and inflammation on the gut microbiome. Using shotgun metagenomic sequencing, we found that microbiome diversity and composition were stable throughout the study duration in both cohorts. Furthermore, we discovered significant differences in the microbiomes between MPN subtypes, such as increased beta-dispersion in subjects with myelofibrosis. Lastly, we found several significant correlations between the abundances of multiple bacterial taxa and cytokine levels. Together, this study provides insight into the interaction between diet, inflammation, and the gut microbiome.

**Importance:** The gut microbiome serves as an interface between the host and diet. Diet and the gut microbiome both play important roles in managing inflammation, which is a key aspect of MPN. Studies have shown that a Mediterranean diet can reduce inflammation. Therefore, we longitudinally characterized the gut microbiomes of MPN patients in response to Mediterranean or US-style dietary counseling to determine whether there were microbiome-associated changes in inflammation. We did not find significant changes in the gut microbiome associated with diet, but we did find several associations with inflammation. This research paves the way for future studies by identifying potential mechanistic targets implicated in inflammation within the MPN gut microbiome.

## Introduction

Myeloproliferative neoplasms (MPN) are a group of hematological malignancies defined by somatic mutations which activate JAK/STAT signaling in hematopoietic stem cells. ^1,2^ This results in an overproduction of myeloid lineage cells. Clinically, MPNs are divided into three clinical phenotypes: polycythemia vera (PV), essential thrombocythemia (ET), and myelofibrosis (MF). PV is characterized by an elevated red blood cell mass. Elevations in platelets and white blood cells are also common. Subjects with ET have elevated platelets but rarely have increased red or white blood cells. MF is characterized by reticulin fibrosis in the bone marrow, and often cytopenia. MF can develop from a “burn out” phase following PV or ET, termed post-PV or post-ET MF, or without a preceding diagnosis of PV or ET, termed primary myelofibrosis.

One feature of MPN is increased inflammatory cytokine abundance, which correlates with worsened symptom burden and disease prognosis. ^3,4^ MPN symptom burden can be severe, and many individuals experience fatigue, early satiety, abdominal discomfort, night sweats, pruritus, bone pain, fever, and unintentional weight loss. Other than bone marrow transplantation, there is no cure for MPN, and management focuses on reducing thrombotic risk and alleviating symptom burden. Current pharmacological treatments for MPN include JAK inhibitors, such as ruxolitinib, but these often carry significant side effects, like immunosuppression. ^5^ Consequently, there is a need to explore low-risk alternatives for MPN management.

One method to non-pharmacologically manage MPN is through the consumption of a Mediterranean (MED) diet, which emphasizes the intake of extra virgin olive oil, fruits, vegetables, whole grains, legumes, fish, nuts, and seeds. A MED diet has been shown to reduce inflammation by lowering C reactive protein and IL-6 levels and is associated with reduced obesity, cardiovascular disease, and cancer risk. ^6–9^ Adherence to a MED diet has been found to alter the gut microbiome, which is the collection of bacteria, fungus, viruses, and other microorganisms living within the large intestine. ^10–15^ Mechanistically, the dietary fiber and unsaturated fat in the MED diet are fermented by gut microbes, resulting in the production of anti-inflammatory metabolites. ^16^ However, it remains to be seen whether a MED diet can be strategically used to manipulate the gut microbiota to promote health by reducing inflammation in MPN.

We performed a randomized clinical trial to investigate whether registered dietician counseling of individuals with MPN can alter their eating patterns toward a Mediterranean style. Subjects were randomly assigned to a (1) MED diet counseling supplemented with complementary extra virgin olive oil (MED cohort) or (2) diet counseling following the standard US Guidelines for Americans (USDA cohort) supplemented with grocery certificates. The study length was 15 weeks, consisting of a 2-week pre-intervention observation, 10 weeks of active dietary counseling, and 3 weeks of post-counseling follow-up. As a key exploratory endpoint, we investigated if a MED diet could produce a microbiome-mediated reduction in inflammation. Blood and stool samples were collected to measure cytokine levels and assess gut microbiome composition, respectively. Survey data was collected to assess the feasibility of a MED diet intervention among MPN patients and symptom burden was tracked using the MPN Symptom Assessment Form (MPN-SAF). In a companion manuscript we describe the relationship between MED diet adherence, symptom burden, and cytokine concentrations. In this manuscript, we detail the association of the gut microbiota with diet, MPN subtype, and cytokine concentrations.

## Results

### Cohort description and study synopsis

Twenty-eight subjects with MPN were recruited for this study (Figure 1). The MED cohort had 15 individuals, while the USDA cohort had 13 individuals. Within the MED cohort, 3 subjects had ET, 4 had MF, and 8 had PV. Within the USDA cohort, 3 subjects had ET, 4 had MF, and 6 had PV. The median age for the MED cohort was 59 +/-14.5 (σ) years, while the median age for the USDA cohort was 61 +/-14 (σ) years. Both groups had 10 females each, with 5 and 3 males in the MED and USDA cohorts, respectively. The study took place over 15 weeks and had an active intervention period from weeks 3-12. Baseline blood and stool samples were collected at week 1, followed by additional sampling during the active intervention at weeks 6 and 9. Follow-up samples were also taken after the intervention’s end at week 15. Throughout the study, six unannounced surveys and 24-hour food recalls (ASA24) were collected to measure diet compliance, and symptom burden was assessed using the MPN-Symptom Assessment Form (MPN-SAF), which grades the 10 most clinically relevant symptoms of MPN patients. ^57^ Table 1 provides a detailed description of each subject’s characteristics.

**Figure 1:**
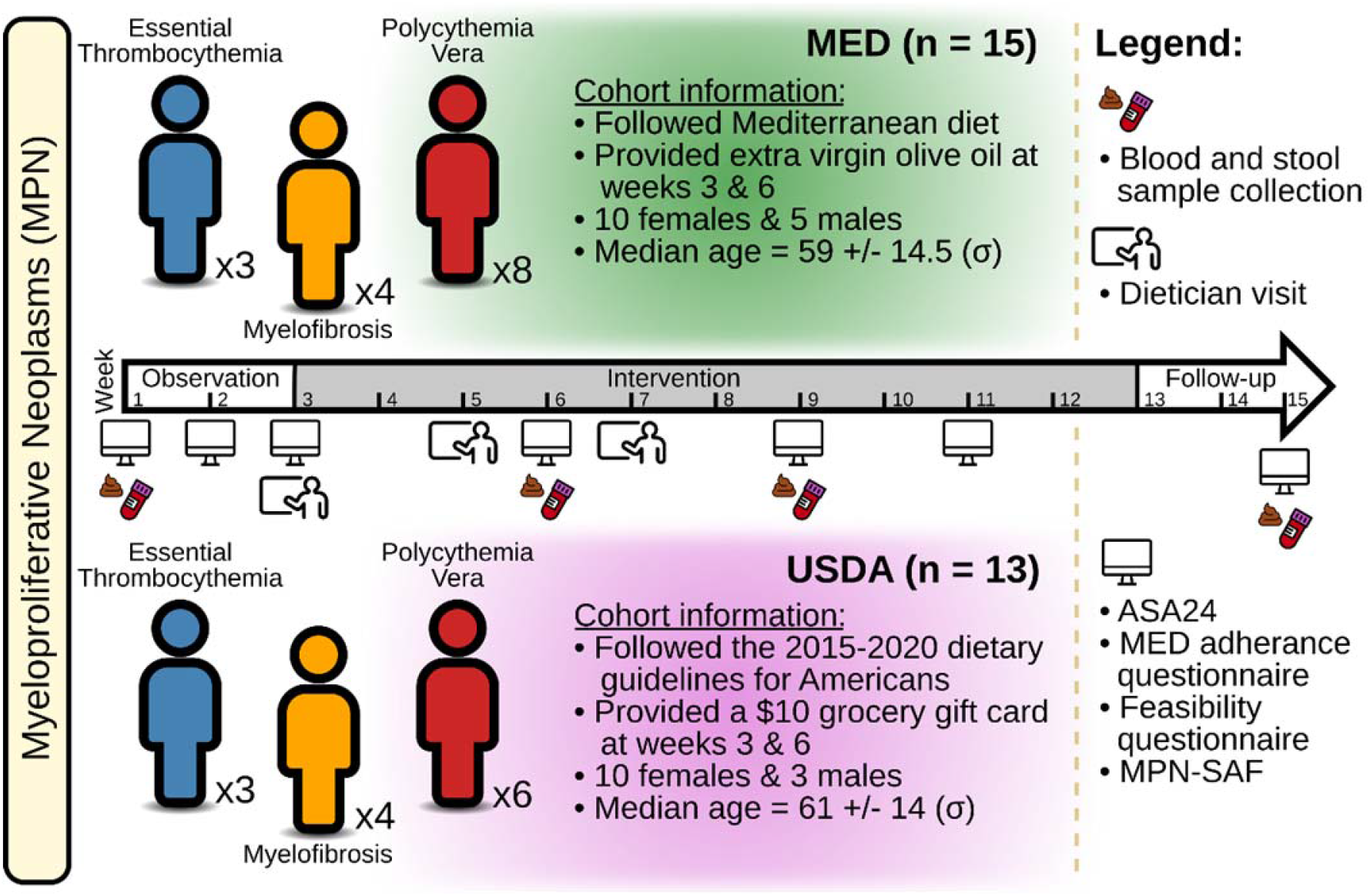
Study design. A total of 28 individuals with myeloproliferative neoplasms (MPN) were enrolled in the study. Participants were randomly assigned to dietary counseling following either a Mediterranean diet (MED, n = 15) or a conventional American diet (USDA, n = 13). The study was 15 weeks long, and had a 2-week observation period, a 10-week intervention period, and a 3-week follow-up period. Blood and stool samples were collected at weeks 1, 6, 9, and 15. At weeks 3, 5, and 7, participants met with a dietician and were informed about the core components of each diet and how to follow it. On weeks 1, 2, 3, 6, 9, 11, and 15, subjects were asked for fill out 24-hour dietary recalls (ASA24), MED adherence and feasibility questionnaires, and an MPN symptom burden assessment (MPN-SAF).

**Table 1:** Subject characteristics. A table detailing each subject’s assigned diet, MPN subtype, treatment, mutation, age, and sex.

| Subject | Diet | MPN | Treatment | Mutation | Age | Sex |
| --- | --- | --- | --- | --- | --- | --- |
| 2 | USDA | PV | Ruxolitinib (Jakafi) | JAK2 | 71 | M |
| 3 | USDA | MF | Observation Only | MPL | 63 | F |
| 5 | USDA | MF | Hydroxyurea (Hydrea), Interferon (Pegasys) | JAK2 | 63 | F |
| 7 | USDA | PV | Interferon (Pegasys) | JAK2 | 44 | F |
| 9 | USDA | ET | Observation Only, Aspirin | JAK2 | 57 | M |
| 10 | USDA | PV | Observation Only, Phlebotomy | JAK2 | 21 | F |
| 12 | USDA | PV | Hydroxyurea (Hydrea) | JAK2 | 77 | M |
| 14 | MED | PV | Aspirin, Interferon (Pegasys) | JAK2 | 34 | F |
| 15 | MED | PV | Hydroxyurea (Hydrea) | JAK2 | 68 | F |
| 16 | MED | ET | Aspirin, Hydroxyurea (Hydrea), Interferon (Pegasys) | JAK2 | 70 | F |
| 17 | MED | PV | Eliquis And Phlebotomy | JAK2 | 58 | F |
| 18 | MED | PV | Hydroxyurea (Hydrea), Prednisone | JAK2 | 66 | M |
| 19 | USDA | ET | Aspirin | JAK2 | 61 | F |
| 20 | MED | ET | Aspirin, Hydroxyurea (Hydrea) | CALR | 71 | M |
| 21 | MED | MF | Aspirin, Hydroxyurea (Hydrea) | CALR | 25 | F |
| 22 | MED | MF | Observation Only, Chinese Herbs | JAK2 | 71 | F |
| 23 | MED | PV | Anagrelide (Agrylin), Ruxolitinib (Jakafi) | JAK2 | 54 | F |
| 24 | MED | MF | Ruxolitinib (Jakafi) | JAK2 | 67 | F |
| 25 | MED | PV | Observation Only, Aspirin, Other | JAK2 | 59 | F |
| 26 | MED | MF | Observation Only, Aspirin | JAK2 | 53 | M |
| 28 | MED | PV | Aspirin, Interferon (Pegasys) | JAK2 | 40 | M |
| 29 | MED | ET | Aspirin, Hydroxyurea (Hydrea) | JAK2 | 70 | F |
| 30 | MED | PV | Observation Only, Aspirin | JAK2 | 50 | M |
| 31 | USDA | ET | Aspirin, Hydroxyurea (Hydrea) | JAK2 | 66 | F |
| 32 | USDA | PV | Aspirin, Hydroxyurea (Hydrea), Other | JAK2 | 67 | F |
| 33 | USDA | PV | Aspirin, Hydroxyurea (Hydrea) | JAK2 | 57 | F |
| 34 | USDA | MF | Other | JAK2 | 51 | F |
| 35 | USDA | MF | Observation Only | JAK2 | 58 | F |

### Gut microbiome diversity and composition is stable during Mediterranean diet intervention

We began our investigation by examining how a Mediterranean Diet (MED) impacts gut microbiome diversity. Analysis of species richness estimates using a linear mixed-effects model (LME) demonstrated that USDA and MED groups did not significantly differ over time after accounting for pre-intervention differences (LME p-value = 0.48, Figure 2A). Analyses of species evenness estimates also showed no differences between diet groups (LME p-value = 0.65, Figure 2B). Sub-setting species richness and evenness comparisons to include only samples from participants highly adherent to a Mediterranean style eating pattern and those least adherent to a Mediterranean style eating pattern during the intervention also did not reveal significant differences (LME richness p-value = 0.48, LME evenness p-value = 0.73).

**Figure 2:**
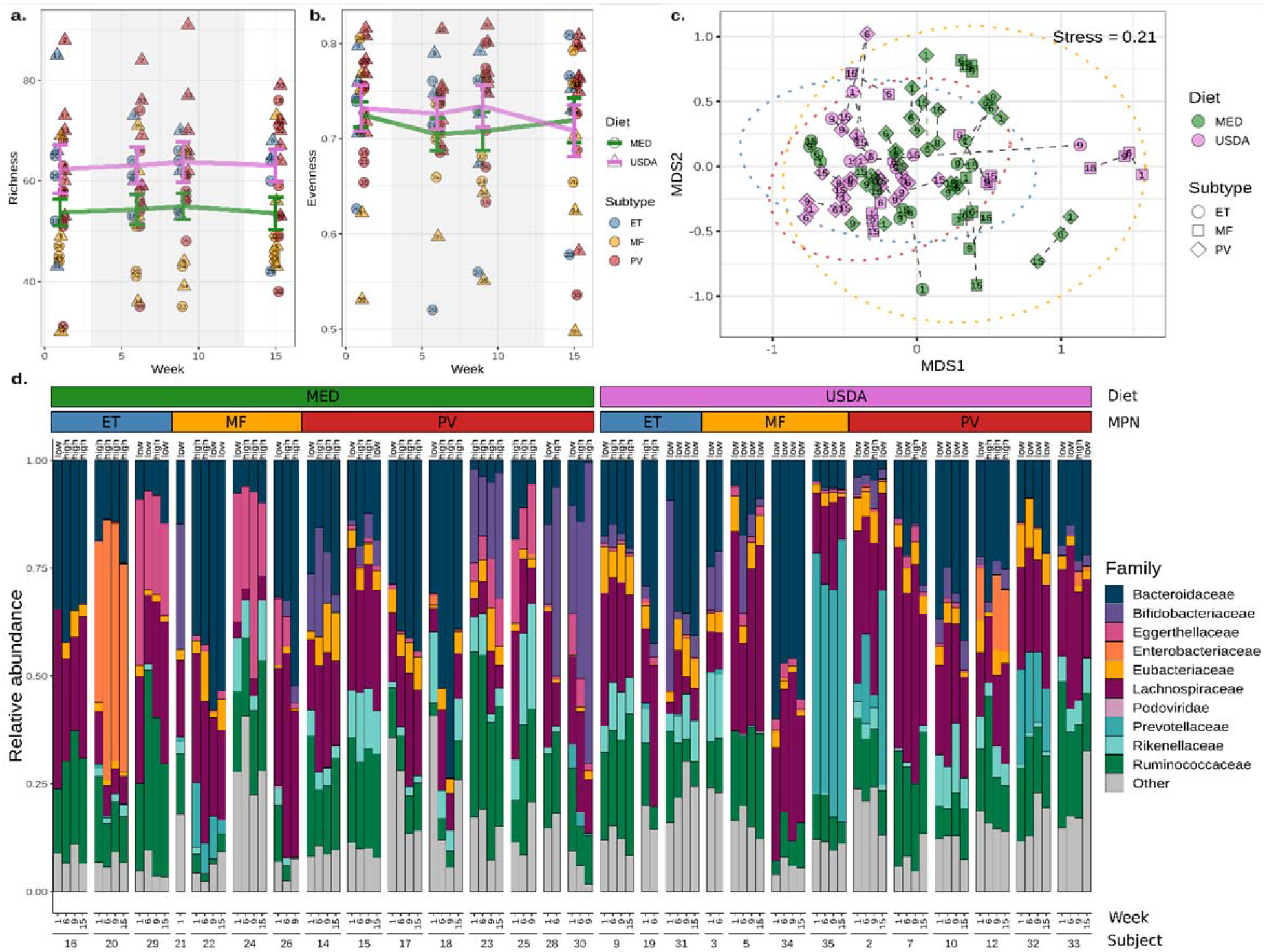
Gut microbiome diversity and composition is stable during Mediterranean diet intervention. **A)** Microbial richness and **B)** evenness estimates of fecal samples collected at weeks 1, 6, 9, and 15. The shaded background indicates the active dietary intervention period for both diet groups. The mean richness or evenness for each group is represented with a colored line, with the error bars reflecting the standard error. Each point is labeled centrally with the individual of origin. **C)** Non-metric multidimensional scaling of Bray-Curtis dissimilarities produced from compositional microbiome data. Points are colored by diet and shaped by MPN subtype. A 95% confidence interval was drawn around each MPN subtype (Blue = ET, Yellow = MF, and Red = PV). Dashed lines connect samples taken from the same individual, and the week of collection is labeled centrally within each point. **D)** A taxa bar plot of the top ten most abundant microbial families across individuals, time, diet, and MPN subtypes. Each sample is labeled ‘high’ or ‘low’ and refers to MED adherence for the week.

Next, we examined the microbial composition, or beta-diversity, of the fecal samples. Species composition analysis using non-metric multidimensional scaling (NMDS) and permutational multivariate analysis of variance (PERMANOVA) showed that there were significant differences associated with MED and USDA groups pre-intervention (Figure 2C; PERMANOVA R^2^ = 0.057, p-value = 0.046, Supplementary Table 1). Therefore, we stratified our PERMANOVA analysis to investigate whether gut microbiome composition changed over time within each individual. This produced non-significant results, suggesting that microbiome composition was stable over the duration of the study (Figure 2D; PERMANOVA R^2^ = 0.007, p-value = 0.76, Supplementary Table 2). Next, PERMANOVA was performed on each MPN subtype to examine whether a specific subtype responded to the diet intervention more than others. No changes were detected in ET (PERMANOVA R^2^ = 0.046, p-value = 0.63), MF (PERMANOVA R^2^ = 0.026, p-value = 0.60), or PV (PERMANOVA R^2^ = 0.016, p-value = 0.77) subtypes over time (Supplementary Table 3). Consequently, we did not find any differentially abundant microbes between MED and USDA groups after adjusting for pre-existing compositional differences.

Characterization of the functional metagenome demonstrated no significant differences between diets as measured by microbial gene richness (LME p-value = 0.65, Supplementary Figure 1A) and gene evenness (LME p-value = 0.19, Supplementary Figure 1B) after accounting for pre-intervention differences. PERMANOVA analysis indicated that there were no significant changes over time within each individual (PERMANOVA R^2^ = 0.009, p-value = 0.29, Supplementary Table 4). Thus, no differentially abundant genes were found between MED and USDA groups.

### Individuals with myelofibrosis have reduced microbial diversity and altered composition

Previous research has demonstrated significant differences in the microbiomes associated with healthy individuals and those with MPN. ^17^ Therefore, we characterized the microbiome between PV, ET, and MF subtypes further. Using species richness estimates, we observed a significant reduction in the number of unique microbes when comparing individuals with MF to PV (Linear mixed-effects p-value = 0.028, Supplementary Figure 2A), and a non-significant reduction when comparing MF to ET (Linear mixed-effects p-value = 0.056, Figure 2A). Species abundance distribution, or evenness, was also reduced in MF, but was not significant compared to PV (LME p-value = 0.12) and ET subtypes (LME p-value = 0.47, Figure 2B).

With respect to beta-diversity, samples from MF were more dissimilar from each other, resulting in a trend towards increased beta-dispersion when compared to ET (LME p-value = 0.089) and PV (LME p-value = 0.056, Supplementary Figure 2C and Figure 3A). Conversely, PV and ET samples tended to cluster together (LME p-value = 0.895, Supplementary Figure 2C and Figure 3A). PERMANOVA demonstrated that the individual of origin significantly explained about 54% of the variance observed in the microbiome, while the MPN subtype significantly explained approximately 6.1% of variance (PERMANOVA p-value = 0.001 for both, Supplementary Table 5). Examining the differential abundance of microbes between MPN subtypes revealed that *Faecalibacterium prausnitzii* was depleted in MF subjects when compared to those with PV or ET (ANCOM2 p < 0.05, Figure 3B). Microbes correlated with *F. prausnitzii* abundance included *Ruminococcus torques, Coprococcus catus, Agathobaculum butyriciproducens, Ruminococcus gnavus, Clostridium bolteae*, and *Blautia sp*. CAG-257 (Figure 3C).

**Figure 3:**
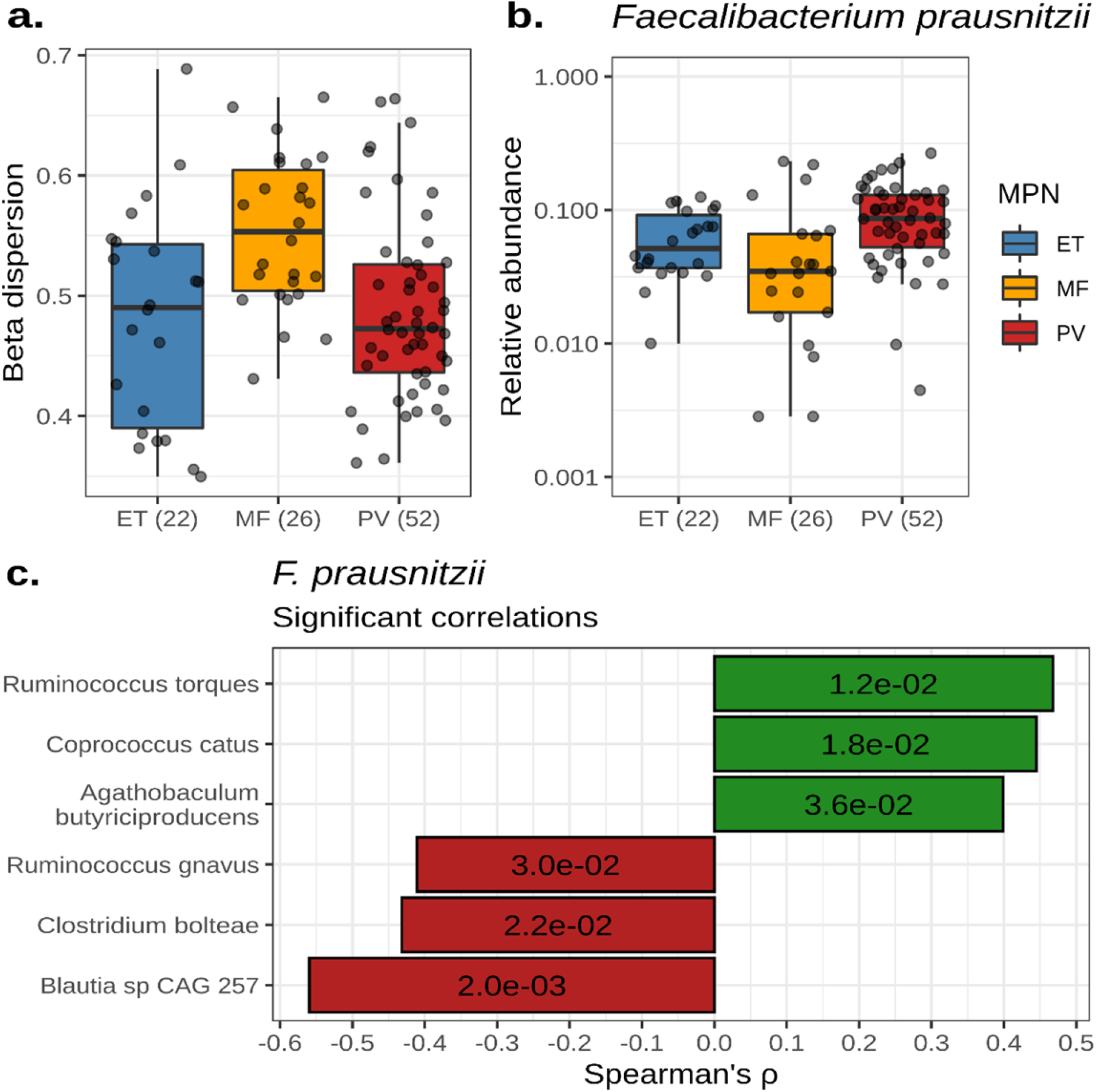
Individuals with myelofibrosis have reduced microbial diversity and altered composition. **A)** A box plot showing the beta-dispersion of each MPN subtype calculated from taxonomic Bray-Curtis dissimilarities. **B)** The relative abundance of *Faecalibacterium prausnitzii* across MPN subtypes. For **A)** and **B)** the number of samples per subtype is labeled parenthetically and the center line within each box defines the median. Boxes define the upper and lower quartiles and whiskers define 1.5x the interquartile range. **C)** A bar plot showing the spearman correlation coefficients of microbes significantly correlated with *F. prausnitzii* abundance. P-values for each correlation are labeled within each bar.

Within the functional metagenome data, we detected a significant reduction in the number of unique microbial genes within MF subjects when compared to PV (LME p-value = 0.016, Supplementary Figure 3A), but not ET (Linear mixed-effects p-value = 0.244, Supplementary Figure 3A). There was no significant difference in the gene evenness among MPN subtypes (Supplementary Figure 3B). NMDS ordination demonstrated that the functional metagenome compositions of MF samples tended to be more disparate from each other when compared to PV (LME p-value = 0.34) and ET (LME p-value = 0.19, Supplementary Figure 3C-D). Additionally, MPN subtype significantly explained about 6.7% of the variance observed in functional metagenome composition (PERMANOVA p-value = 0.001, Supplementary Table 6), while the individual of origin was associated with about 54% of the variance (PERMANOVA p-value = 0.001, Supplementary Table 6). Differential abundance analysis produced no significantly different genes between MPN subtypes after FDR correction.

### Cytokine levels are correlated with microbiome diversity and composition

After subsequent analysis of MPN subtypes and their gut microbiomes, we next asked if diversity or composition of the microbiome were associated with the levels of ten cytokines. Comparison of cytokine concentrations between MPN subtypes revealed a significant increase of TNFα and IL-12p70 in subjects with MF when compared to ET (Tukey’s test; TNFα p-adj < 0.001 and IL-12p70 p-adj = 0.016) and PV (Tukey’s test; TNFα p-adj = 0.002 and IL-12p70 p-adj = 0.022, Figure 4A). IL-6, IL-8, and IL-10 concentrations were elevated in subjects with MF but were not statistically significant (Figure 4A). Microbial richness was negatively correlated with TNFα (Spearman’s ρ = −0.50, p-adj = 0.07, Figure 4B) and IL-12p70 (Spearman’s ρ = −0.45, p-adj = 0.15, Figure 4C).

**Figure 4:**
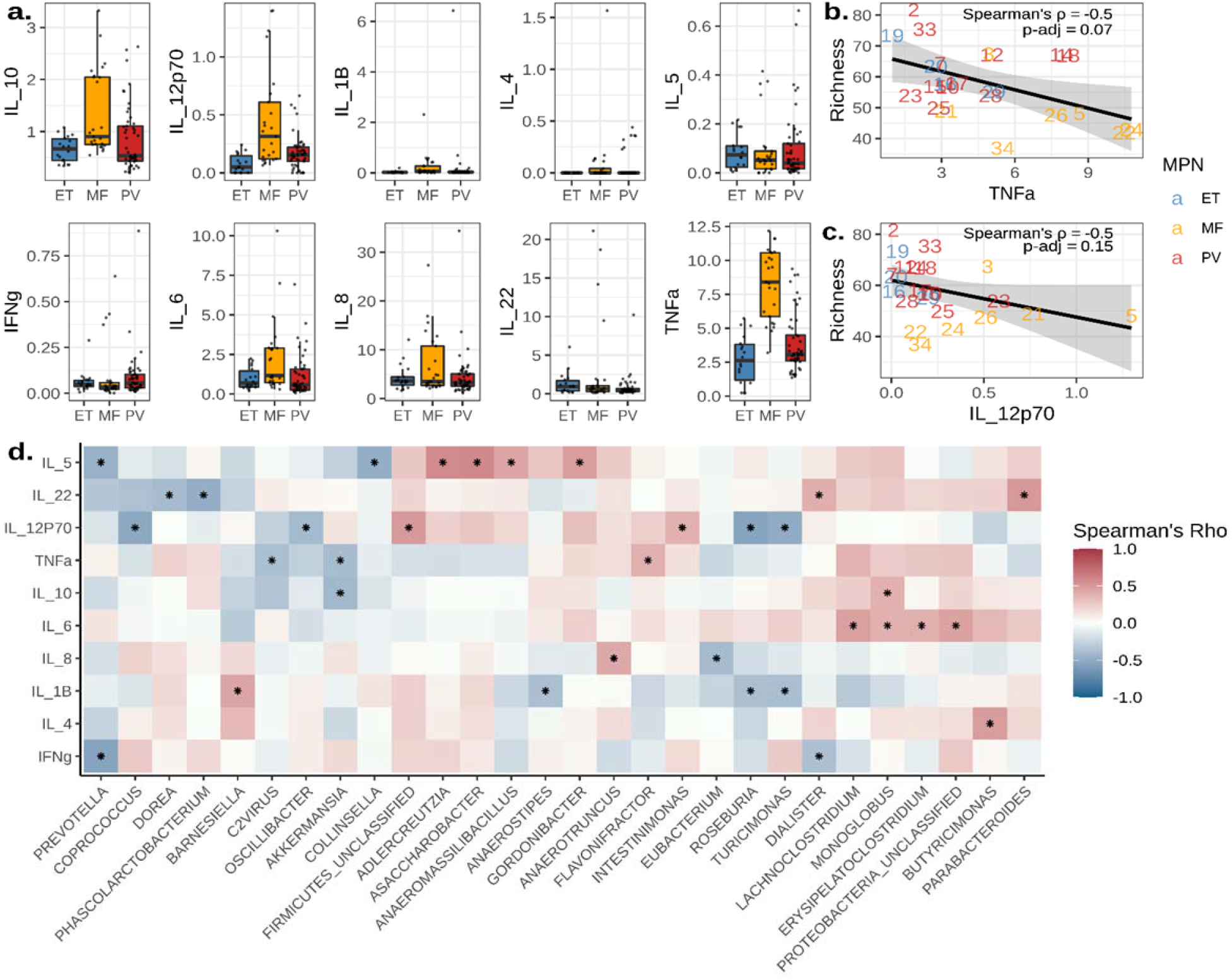
Cytokine levels are correlated with microbiome diversity and composition. **A)** Box plots displaying the concentration of cytokines measured in pg/mL across MPN subtypes. The center line within each box defines the median, boxes define the upper and lower quartiles, and whiskers define 1.5x the interquartile range. **B-C)** Scatter plots of TNFα (**B**) and IL-12p70 (**C**) concentrations in pg/mL correlated with species richness estimates. Points are labeled by the individual of origin and colored by MPN subtypes. A line represents the mean, and the shaded area delineates the 95% confidence interval. **D)** A heat map of microbial genera whose abundances significantly correlated with cytokine concentrations. Asterisks denote significant correlations (p < 0.05).

Next, we correlated cytokine levels with microbial abundances at the genus level, resulting in 34 significant correlations (Figure 4D). Notable correlations included associations with TNFα vs. *Flavonfractor* (Spearman’s ρ = 0.39, p = 0.038), IL-12p70 vs. *Roseburia* (Spearman’s ρ = −0.55, p = 0.002), and IL-8 vs. *Eubacterium* (Spearman’s ρ = −0.41, p = 0.032, Supplementary Figure 4). Similarly, we compared cytokine levels with functional pathway abundances, producing 162 significant correlations (Supplementary Figure 5). Notable correlations included TNFα vs. 4-deoxy-L-threo-hex-4-enopyranuronate degradation (Spearman’s ρ = −0.42, p = 0.038), TNFα vs. β-(1,4)-mannan degradation (Spearman’s ρ = −0.48, p = 0.019), and IL-12p70 vs. GDP-mannose biosynthesis (Spearman’s ρ = −0.59, p = 0.003, Supplementary Figure 6).

## Discussion

Our goal with this manuscript was to 1) assess whether a MED diet altered the gut microbiome of subjects with MPN and 2) to investigate the association between diversity and composition of the gut microbiome and levels of several cytokines. In a separate manuscript, we describe the feasibility of a MED diet intervention in the MPN subject population, changes in macronutrients associated with the dietary intervention, and the interaction between diet adherence, symptom burden, and cytokine concentrations. Here, we report that microbial diversity or composition did not significantly change during a MED diet intervention over a 10-week active dietary intervention period. Instead, we found the MPN subtype played a greater role in determining microbiome diversity and composition. Individuals with ET and PV had more similar microbial compositions, while those with MF were more disparate. Furthermore, a reduction of microbial diversity correlated with elevated TNFα and IL-12p70 concentrations in subjects with MF. These differences in cytokine concentrations were associated with the abundances of 34 microbial genera and 162 metabolic pathways, further establishing a role for the gut microbiome in inflammation and MPN.

With respect to diet-mediated changes in the microbiome, there are multiple explanations as to why the microbiomes of individuals remained stable throughout the dietary intervention. The first is intervention duration. Long term adherence to a Mediterranean diet has been demonstrated to reduce the incidence of cardiovascular disease, Alzheimer’s disease, colorectal cancer, diabetes, and obesity. ^8,9^ Studies performing MED diet interventions have ranged from 6 weeks to 7 years. ^12–15,18–26^ Of studies examining gut microbiome composition, the 6-week MED diet intervention performed by Marlow *et al* yielded no significant differences in gut microbiome composition or CRP levels in subjects with Crohn’s disease. ^26^ Comparatively, Nagpal *et al* conducted a MED diet intervention in a non-human primate model over the course of 2.5 years, where a significant difference in the microbiome was observed between macaques who consumed a Western diet versus a MED diet. ^14^ Although diet has been shown to rapidly alter the composition of the microbiome, our study and others suggest that longer dietary interventions are needed to detect changes in gut microbiome composition, especially with smaller sample sizes. ^27^

Another consideration in the successful manipulation of gut microbiomes with diet is presence of specific microbial taxa, functions, or enterotypes. Stratification of microbiomes into enterotypes has revealed enterotype-specific predictors to dietary intervention response. ^28^ Klimenko *et al*. found that the strongest predictor of whether an individual would respond to a dietary intervention was the average number of genes per microbe. ^28^ A negative correlation between the average number of genes per microbe and alpha diversity was found, suggesting that more diverse communities are formed by specialist microbes with fewer genes. ^28^ The microbiomes associated with industrialized countries, like the United States, often have reduced diversity and a higher abundance of *Bacteroides* when compared to non-industrialized countries. ^29^ Many *Bacteroides* are generalists, meaning they contain more genes and wider metabolic potentials than specialist taxa. ^30^ The predominance of generalist taxa has been known to contribute to microbiome stability. ^31^ Therefore, it is plausible that the microbiomes of industrialized individuals have evolved to resist perturbations, such as those caused by antibiotic usage or short-term dietary changes. In one study, individuals with a higher ratio of *Prevotella* to *Bacteroides* lost more weight than those with a lower *Prevotella* to *Bacteroides* ratio while consuming a New Nordic Diet, suggesting that a higher abundance of generalists is associated with intervention outcome too. ^32^ Our samples contained relatively high abundances of *Bacteroides*, so it is possible that the microbiomes of these individuals were resistant to short-term dietary changes as reflected by the non-significant changes in diversity, composition, and function over time.

One final factor which could affect the stability of microbiomes is the strength of the dietary intervention. Due to differences in agriculture and food processing, a MED diet in United States is likely different than a MED diet in the Mediterranean region. This can affect the number of antibiotic and prebiotic compounds found in each diet. One prebiotic component of the MED diet that can influence gut microbiome composition is extra virgin olive oil (EVOO). EVOO is rich in polyphenols and oleic acid, which have been demonstrated to have anti-oxidative and anti-inflammatory properties. ^33,34^ Over 90% of polyphenols are digested and metabolized in the colon by the gut microbiota. ^35^ Dietary supplementation of EVOO in humans has been shown to promote the growth of beneficial microbes like *Bifidobacterium* and lactic acid producing bacteria. ^33,36^ In rodent models, consumption of EVOO results in an increased abundance of *Bifidobacterium, Lactobacillus*, and *Clostridium*. ^37,38^

The MED diet is also typically higher in dietary fiber when compared to a typical USDA diet. Dietary fiber is fermented by the gut microbiota to produce short-chain fatty acids, like acetate, propionate, and butyrate. Butyrate is critical for gut health, as it the primary source of energy for colonocytes and reduces inflammation by stimulating the production of T-regulatory cells and IL-10 producing cells. ^39^ In this study, we did not find that the abundances of butyrate-producing bacteria differed between diets. Instead, we saw a reduction of the butyrate-producing microbe, *F. prausnitzii*, in subjects with MF. We also noted significant positive correlations between *F. prausnitzii, Agathobaculum butyriciproducens*, and *Coprococcus catus* abundances. *A. butyriciproducens* is a butyrate-producing microbe, while *C. catus* produces butyrate and propionate. ^40^ We also observed broader, community wide differences between subjects with ET, PV, and MF. Notably, the microbiome composition of MF subjects was more dissimilar to each other when compared to ET and PV. Our previous work comparing the gut microbiome composition of healthy and MPN subjects similarly showed that individuals with MF had increased beta dispersion when compared to ET and PV. ^17^ These results describe a phenomenon known as the ‘Anna Karenina principal’ for animal microbiomes, which states that stressors affect microbiomes in unpredictable ways, leading to increased community beta-dispersion. ^41,42^

A likely stressor resulting in higher MF beta-dispersion is the increased concentration of pro-inflammatory cytokines. Inflammation has been known negatively affect the gut microbiome. Supporting this notion, we found that TNFα and IL-12p70 were significantly increased in MF subjects, which negatively correlated with species richness overall. We found IL-12p70 negatively correlated with the genus *Roseburia*, which are butyrate-producing microorganisms known to alleviate inflammation by promoting T-regulatory cell differentiation. ^43,44^ We also observed a significant negative correlation with *Eubacterium* and the pro-inflammatory cytokine, IL-8. *Eubacterium* also produce butyrate and have been shown to lessen inflammation by promoting IL-10 production. ^43,45^ TNFα levels and the abundance of the 4-deoxy-L-threo-hex-4-enopyranuronate degradation pathway were negatively correlated as well. This pathway plays a role in the degradation of uronic acids, such as apple pectin. β-(1,4)-mannan is another compound found in plant cell walls and the pathway abundance for its degradation was found to be negatively correlated with TNFα levels.

Taken together, it is possible that the increased inflammation observed in individuals with MPN, particularly MF, is exacerbated by the lack of sufficient short-chain fatty acid production. The MED diet has been previously shown to promote the growth of *F. prausnitzii* specifically, therefore, future experiments could attempt to restore microbial short-chain fatty acid production to reduce inflammation. When designing dietary interventions, however, special attention should be given to the intervention duration and the ability for existing gut microbes to use and respond to prebiotic compounds. This may ensure that the desired outcomes are achieved, allowing us to manipulate the gut microbiome to promote health and ameliorate disease.

## Methods

### Recruitment of subjects

Patients were recruited between October 2018 and September 2019. Participants were included if they were over the age of 18 with a previous diagnosis of a Philadelphia chromosome negative MPN (including PV, ET, MF), had an ECOG score of 2 or less, a life expectancy of greater than 20 weeks, had internet access, an email address, and could read and understand English. Any type of previous or current therapy was also allowed. Participants were excluded if they were pregnant or planning on becoming pregnant, lost more than 10 pounds or 10% of their body weight in the last 6 months, or were allergic to nuts and olive oil. Forty-seven participants were screened. Five did not meet the inclusion criteria, and an additional 11 subjects were excluded due to incomplete survey data during the observation period. Thirty-one subjects were randomly assigned to a diet, but 2 withdrew participation and one failed to provide sufficient survey data. The final number of study participants was 28, with 15 belonging to the MED cohort and 13 belonging to the USDA cohort.

### Collection of dietary intervention feasibility, adherence, and symptom burden data

During the first week of the intervention period, each participant met individually with a dietician to learn about the central components of their assigned diet. There were follow-up dietician visits during weeks 5 and 7. Participants were emailed educational materials on their respective diet weekly during the 10-week active intervention period. Furthermore, participants in the MED cohort were given 750 mL of extra virgin olive oil and those in the USDA cohort were given a $10 USD grocery gift card at weeks 3 and 6. Throughout the study, participants were required to fill out 4 unannounced surveys given during weeks 1, 2, 3, 6, 9, 11, and 15. The first survey measured dietary intervention feasibility and asked, “how easy is it for you to follow this diet, with 1 being very easy to follow and 10 being very difficult to follow?” The second survey measured MED diet adherence. For this, the established 14-item Mediterranean diet adherence score (MEDAS) was used. ^46^ Adherence to a MED diet was defined as a “high” for the week if a score of >8/14 was obtained. Next, we asked subjects to complete 24-hour food recalls by using the Automated Self-Administered 24-hour Dietary Assessment Tool (ASA24). Lastly, symptom burden was assessed via the MPN symptom assessment form (MPN-SAF), which grades the 10 most clinically relevant MPN symptoms. ^47^ Surveys were administered through email.

### Blood collection and cytokine measurements

Peripheral blood was drawn on weeks 1, 3, 6, and 15 in tubes containing ethylenediaminetetraacetic acid (EDTA). Plasma was obtained by centrifuging 3-4 ml of blood for 10 minutes at 2500 rpm, aliquoted, and was stored at −80°C. Frozen plasma was sent to Quanterix in Billerica, MA for analysis. A Human CorPlex 10 Cytokine Array kit #85-0329 (IL-12p70, IL-1B, IL-4, IL-5, IFNγ, IL-6, IL-8, IL-22, TNFα, and IL-10) was used according to manufacturer’s protocol and analyzed using A Quanterix SPX imager system on-site at Quanterix Headquarters in Billerica, MA.

### Fecal sample collection

To perform gut microbiome analysis, four stool samples were requested from each participant over the course of the 15-week trial. The samples were collected by the participants themselves using Zymo RNA/DNA shield fecal collection tubes (Cat. #R1101) during weeks 1, 6, 9, and 15. Samples were returned in person or by mail. In total, 103 samples were collected. Samples were stored at −80°C once returned.

### DNA extraction

Fecal samples stored in DNA/RNA shield were thawed on ice, vortexed to homogenize, then DNA from 1000 uL of fecal slurry was extracted using ZymoBiomics DNA Miniprep Kit (Cat. #D4300) according to the manufacturer’s protocol. Bead lysis during the extraction was performed at 6.5 m/s for 5 minutes total (MPBio FastPrep-24).

### Shotgun library preparation and sequencing

Libraries for shotgun sequencing were prepared using the Illumina DNA prep kit (Cat. # 20018705), using an adapted low-volume protocol. ^48^ In summary, we reduced the amount of DNA used per sample to a maximum of 5 uL or 50 ng (whichever was reached first). Tagmentation was performed according to the manufacturer’s protocol, but volumes were reduced to 1 uL of bead-linked transposome and tagmentation buffer each. Next, 1.25 uL of 1 uM i5 and i7 indices were added to each sample and annealed via polymerase chain reaction using 10 uL of KAPA HiFi HotStart ReadyMix (Cat. # 7958935001). Afterwards, libraries were combined, size-selected, and cleaned using 56 and 14.4 uL of sample purification beads according to the low-volume protocol. Positive and negative sequencing controls were included during the library preparation using the ZymoBIOMICS Microbial Community DNA Standard (Cat. #D6305) and purified water, respectively. The quality of libraries was assessed with Quanti-iT PicoGreen dsDNA (Cat. #P7589) for quantity and Agilent Bioanalyzer High Sensitivity DNA Analysis (Cat. #5067-4626) for fragment size. Libraries were shipped overnight on dry ice to Novogene Corporation Inc. (Sacramento, CA) to be sequenced using Illumina’s Hiseq 4000. An average of 2,819,107 +/-670,543 (σ) paired-end reads per sample, 150 basepairs long, were obtained.

### OTU table generation

Raw data was first cleaned to remove sequencing adapters and artifacts using the BBMap v38.79 script ‘bbduk.sh’ with the flag ‘ref=adapters,artifacts’. ^49^ BBMap’s ‘demuxbyname.sh’ was used to demultiplex sequences using the default parameters. Quality filtering of sequences was performed using PRINSEQ++ v1.2 with the following parameters: -trim_left 5 -trim_right 5 -min_len 100 -trim_qual_right 28 -min_qual_mean 25. ^50^ Quality checking was done with FastQC v0.11.8 on default parameters. This resulted a mean and standard deviation of 2,731,886 +/-648,042 paired-end reads, respectively. Human-derived reads were removed using BowTie2 v2.4.5 using the default parameters and hg38 as the reference genome, which produced an average of 2,498,159 +/-960,477 (σ) reads per sample. ^51^ Taxonomic assignment of sequence data was performed using MetaPhlAn v3.0.14 with the default parameters and the CHOCOPhlAn v2019.01 database. ^52^

### Microbiome functional potential data generation

Individual gene annotations were produced by first cross-assembling reads into contiguous sequences using MEGAHIT v1.1.1 with a minimum length of 2,500 base pairs and the flag ‘-k-list 31,41,51,61,71,81,91,101,111’. ^53^ Afterwards, open reading frames were assigned with Prodigal v2.6.3 and then annotated with eggNOG mapper v2.0 using the eggNOG v5.0 database. ^54,55^ Next, BowTie2 v2.4.5 was used to align samples to the annotated genes to obtain a table of per sample counts for each gene. Lastly, per sample gene counts were normalized to reads per kilobase per genome equivalent using MicrobeCensus v1.1.1 on default parameters. ^56^ For the functional annotation of metabolic pathways, we ran our quality-filtered, unassembled reads through HUMAnN v3.0.1 using the default parameters and the UniRef90 v201901b database. ^52^ The ‘humann_renorm_table’ and ‘humann_join_tables’ scripts were used to create a pathway abundance table of normalized counts in copies per million.

### Data analysis

Data analysis of OTUs, genes, and pathways was performed in R v4.2.1. The first step was removing microbes or genes which contaminated our sequencing controls from all samples. The Vegan v2.6-2 package was used to calculate the following metrics: richness with the ‘specnumber’ function, evenness with the formula ‘diversity(x, index = “Shannon”) / log_10_(specnumber(x))’, Bray-Curtis beta diversity with the ‘vegdist’ function, PERMANOVA with the ‘adonis2’ function, NMDS with the ‘metaMDS’ function, and beta-dispersion with the ‘betadisper’ function. Please see Supplementary Tables 1 – 6 for PERMANOVA formulas and parameters. Significance testing of richness, evenness, and beta dispersion was performed using linear-mixed effect models with the nlme v3.1-159 package. Significance testing of cytokine concentrations was done using an ANOVA and Tukey’s post-hoc test with the ‘aov’ and ‘TukeyHSD’ functions. Spearman correlations were obtained using the ‘cor.test’ and ‘rcorr’ functions. Differential abundance of OTUs was determined with ANCOM v2.1 with the parameters: rand_formula = “~ 1 | Subject”, p_adj_method = “none”, alpha = 0.05. We were unable to perform ANCOM for gene and pathway abundances, therefore, we averaged abundances within each subject to eliminate repeated measurements and performed a Kruskal-Wallis test. When appropriate, multiple comparisons were corrected for using the ‘p.adjust(x, method = “fdr”)’ function. All code, scripts, and parameters for data processing and analysis can be found at https://github.com/Javelarb/MPN_diet_intervention.

## Declarations

### Data availability statement

Sequencing data is available on the Sequence Read Archive under the BioProject ID, PRJNA918651. The taxonomic table with corresponding metadata used to generate the source data can be found in ‘Supplementary_file_2.csv’. Larger files used in our functional metagenomic analysis are available on the Dryad Digital Repository (Reviewer Link). Additional data and materials are available upon reasonable request.

### Code availability statement

All code for data processing and analysis are available on GitHub at: https://github.com/Javelarb/MPN_diet_intervention.

## Acknowledgements

This study was partially funded by the UCI Anti-Cancer Challenge Grant.

## Author contributions

AGF developed the study, recruited, and consented patients, and analyzed data, LFML recruited and consented patients, collected and analyzed data, JN coordinated patient visits and collected clinical data, HN coordinated patient visits and collected clinical data, AO developed the study and analyzed data, JAB conducted a portion of the experiments and performed the data analysis. JAB wrote the manuscript with guidance from AF and KW.

## Ethics approval

This study was approved by the Institutional Review Board (IRB) of the University of California, Irvine (HS#2018-4521) and registered on clinicaltrials.gov (NCT03907436).

## References

1. Kralovics, R. et al. A Gain-of-Function Mutation of JAK2 in Myeloproliferative Disorders. N Engl J Med 352, 1779–1790 (2005).

2. Nangalia, J. et al. Somatic CALR Mutations in Myeloproliferative Neoplasms with Nonmutated JAK2. N Engl J Med 369, 2391–2405 (2013).

3. Fleischman, A. G. Inflammation as a Driver of Clonal Evolution in Myeloproliferative Neoplasm. Mediators of Inflammation 2015, 1–6 (2015).

4. Craver, B., El Alaoui, K., Scherber, R. & Fleischman, A. The Critical Role of Inflammation in the Pathogenesis and Progression of Myeloid Malignancies. Cancers 10, 104 (2018).

5. Tefferi, A. JAK inhibitors for myeloproliferative neoplasms: clarifying facts from myths. Blood 119, 2721–2730 (2012).

6. Smidowicz, A. & Regula, J. Effect of Nutritional Status and Dietary Patterns on Human Serum C-Reactive Protein and Interleukin-6 Concentrations. Advances in Nutrition 6, 738–747 (2015).

7. Estruch, R. Anti-inflammatory effects of the Mediterranean diet: the experience of the PREDIMED study. Proc. Nutr. Soc. 69, 333–340 (2010).

8. Sofi, F., Abbate, R., Gensini, G. F. & Casini, A. Accruing evidence on benefits of adherence to the Mediterranean diet on health: an updated systematic review and meta-analysis. The American Journal of Clinical Nutrition 92, 1189–1196 (2010).

9. Gotsis, E. et al. Health Benefits of the Mediterranean Diet: An Update of Research Over the Last 5 Years. Angiology 66, 304–318 (2015).

10. De Filippis, F. et al. High-level adherence to a Mediterranean diet beneficially impacts the gut microbiota and associated metabolome. Gut 65, 1812–1821 (2016).

11. Mitsou, E. K. et al. Adherence to the Mediterranean diet is associated with the gut microbiota pattern and gastrointestinal characteristics in an adult population. Br J Nutr 117, 1645–1655 (2017).

12. Ghosh, T. S. et al. Mediterranean diet intervention alters the gut microbiome in older people reducing frailty and improving health status: the NU-AGE 1-year dietary intervention across five European countries. Gut 69, 1218–1228 (2020).

13. Meslier, V. et al. Mediterranean diet intervention in overweight and obese subjects lowers plasma cholesterol and causes changes in the gut microbiome and metabolome independently of energy intake. Gut 69, 1258–1268 (2020).

14. Nagpal, R. et al. Gut Microbiome Composition in Non-human Primates Consuming a Western or Mediterranean Diet. Front. Nutr. 5, 28 (2018).

15. Haro, C. et al. Two Healthy Diets Modulate Gut Microbial Community Improving Insulin Sensitivity in a Human Obese Population. The Journal of Clinical Endocrinology & Metabolism 101, 233–242 (2016).

16. Bailey, M. A. & Holscher, H. D. Microbiome-Mediated Effects of the Mediterranean Diet on Inflammation. Advances in Nutrition 9, 193–206 (2018).

17. Oliver, A. et al. Fecal Microbial Community Composition in Myeloproliferative Neoplasm Patients Is Associated with an Inflammatory State. Microbiol Spectr 10, e00032–22 (2022).

18. Shively, C. A. et al. Mediterranean versus Western Diet Effects on Caloric Intake, Obesity, Metabolism, and Hepatosteatosis in Nonhuman Primates: Mediterranean and Western Diet in Nonhuman Primates. Obesity 27, 777–784 (2019).

19. Garcia-Rios, A. et al. Beneficial effect of CLOCK gene polymorphism rs1801260 in combination with low-fat diet on insulin metabolism in the patients with metabolic syndrome. Chronobiology International 31, 401–408 (2014).

20. Kaaks, R. et al. Effects of dietary intervention on IGF-I and IGF-binding proteins, and related alterations in sex steroid metabolism: the Diet and Androgens (DIANA) Randomised Trial. Eur J Clin Nutr 57, 1079–1088 (2003).

21. Delgado-Lista, J. et al. CORonary Diet Intervention with Olive oil and cardiovascular PREVention study (the CORDIOPREV study): Rationale, methods, and baseline characteristics. American Heart Journal 177, 42–50 (2016).

22. Shai, I. et al. Weight Loss with a Low-Carbohydrate, Mediterranean, or Low-Fat Diet. N Engl J Med 359, 229–241 (2008).

23. Mekki, K., Bouzidi-bekada, N., Kaddous, A. & Bouchenak, M. Mediterranean diet improves dyslipidemia and biomarkers in chronic renal failure patients. Food & Funct. 1, 110 (2010).

24. Paniagua, J. A. et al. A MUFA-Rich Diet Improves Posprandial Glucose, Lipid and GLP-1 Responses in Insulin-Resistant Subjects. Journal of the American College of Nutrition 26, 434–444 (2007).

25. Salas-Salvadó, J. et al. Prevention of Diabetes With Mediterranean Diets: A Subgroup Analysis of a Randomized Trial. Ann Intern Med 160, 1–10 (2014).

26. Marlow, G. et al. Transcriptomics to study the effect of a Mediterranean-inspired diet on inflammation in Crohn’s disease patients. Hum Genomics 7, 24 (2013).

27. David, L. A. et al. Diet rapidly and reproducibly alters the human gut microbiome. Nature 505, 559–563 (2014).

28. Klimenko, N. S., Odintsova, V. E., Revel-Muroz, A. & Tyakht, A. V. The hallmarks of dietary intervention-resilient gut microbiome. npj Biofilms Microbiomes 8, 77 (2022).

29. Smits, S. A. et al. Seasonal cycling in the gut microbiome of the Hadza hunter-gatherers of Tanzania. Science 357, 802–806 (2017).

30. Sriswasdi, S., Yang, C. & Iwasaki, W. Generalist species drive microbial dispersion and evolution. Nat Commun 8, 1162 (2017).

31. Matias, M. G., Combe, M., Barbera, C. & Mouquet, N. Ecological strategies shape the insurance potential of biodiversity. Front. Microbio. 3, (2013).

32. Hjorth, M. F. et al. Pre-treatment microbial Prevotella-to-Bacteroides ratio, determines body fat loss success during a 6-month randomized controlled diet intervention. Int J Obes 42, 580–583 (2018).

33. Luisi, M. L. E. et al. Effect of Mediterranean Diet Enriched in High Quality Extra Virgin Olive Oil on Oxidative Stress, Inflammation and Gut Microbiota in Obese and Normal Weight Adult Subjects. Front. Pharmacol. 10, 1366 (2019).

34. Millman, J. et al. Metabolically and immunologically beneficial impact of extra virgin olive and flaxseed oils on composition of gut microbiota in mice. Eur J Nutr 59, 2411–2425 (2020).

35. Ozdal, T. et al. The Reciprocal Interactions between Polyphenols and Gut Microbiota and Effects on Bioaccessibility. Nutrients 8, 78 (2016).

36. Martín-Peláez, S. et al. Effect of virgin olive oil and thyme phenolic compounds on blood lipid profile: implications of human gut microbiota. Eur J Nutr 56, 119–131 (2017).

37. Zhao, Z., Shi, A., Wang, Q. & Zhou, J. High Oleic Acid Peanut Oil and Extra Virgin Olive Oil Supplementation Attenuate Metabolic Syndrome in Rats by Modulating the Gut Microbiota. Nutrients 11, 3005 (2019).

38. Hidalgo, M. et al. Changes in Gut Microbiota Linked to a Reduction in Systolic Blood Pressure in Spontaneously Hypertensive Rats Fed an Extra Virgin Olive Oil-Enriched Diet. Plant Foods Hum Nutr 73, 1–6 (2018).

39. Chen, J. & Vitetta, L. Inflammation-Modulating Effect of Butyrate in the Prevention of Colon Cancer by Dietary Fiber. Clinical Colorectal Cancer 17, e541–e544 (2018).

40. Reichardt, N. et al. Phylogenetic distribution of three pathways for propionate production within the human gut microbiota. ISME J 8, 1323–1335 (2014).

41. Zaneveld, J. R., McMinds, R. & Vega Thurber, R. Stress and stability: applying the Anna Karenina principle to animal microbiomes. Nat Microbiol 2, 17121 (2017).

42. Kaszubinski, S. F. et al. Dysbiosis in the Dead: Human Postmortem Microbiome Beta-Dispersion as an Indicator of Manner and Cause of Death. Front. Microbiol. 11, 555347 (2020).

43. Kumari, M. et al. Fostering next-generation probiotics in human gut by targeted dietary modulation: An emerging perspective. Food Research International 150, 110716 (2021).

44. Zhu, C. et al. Roseburia intestinalis inhibits interleukin-17 excretion and promotes regulatory T cells differentiation in colitis. Mol Med Report (2018) doi:10.3892/mmr.2018.8833.

45. Chung, W. S. F. et al. Prebiotic potential of pectin and pectic oligosaccharides to promote anti-inflammatory commensal bacteria in the human colon. FEMS Microbiology Ecology 93, (2017).

46. Martínez-González, M. A. et al. A 14-Item Mediterranean Diet Assessment Tool and Obesity Indexes among High-Risk Subjects: The PREDIMED Trial. PLoS ONE 7, e43134 (2012).

47. Scherber, R. et al. The Myeloproliferative Neoplasm Symptom Assessment Form (MPN-SAF): International Prospective Validation and Reliability Trial in 402 patients. Blood 118, 401–408 (2011).

48. Weihe, C. & Avelar-Barragan, J. Next generation shotgun library preparation for Illumina sequencing - low volume v1. doi:10.17504/protocols.io.bvv8n69w.

49. Bushnell, B. BBMap: A Fast, Accurate, Splice-Aware Aligner. (2014).

50. Cantu, V. A., Sadural, J. & Edwards, R. PRINSEQ++, a multi-threaded tool for fast and efficient quality control and preprocessing of sequencing datasets. https://peerj.com/preprints/27553v1 (2019) doi:10.7287/peerj.preprints.27553v1.

51. Langmead, B. & Salzberg, S. L. Fast gapped-read alignment with Bowtie 2. Nat Methods 9, 357–359 (2012).

52. Beghini, F. et al. Integrating taxonomic, functional, and strain-level profiling of diverse microbial communities with bioBakery 3. eLife 10, e65088 (2021).

53. Li, D. et al. MEGAHIT v1.0: A fast and scalable metagenome assembler driven by advanced methodologies and community practices. Methods 102, 3–11 (2016).

54. Hyatt, D. et al. Prodigal: prokaryotic gene recognition and translation initiation site identification. BMC Bioinformatics 11, 119 (2010).

55. Huerta-Cepas, J. et al. eggNOG 5.0: a hierarchical, functionally and phylogenetically annotated orthology resource based on 5090 organisms and 2502 viruses. Nucleic Acids Research 47, D309–D314 (2019).

56. Nayfach, S. & Pollard, K. S. Average genome size estimation improves comparative metagenomics and sheds light on the functional ecology of the human microbiome. Genome Biol 16, 51 (2015).

